# Systemic and intra-amygdala administrations of midazolam reverse anxiety-like behavior induced by cohabiting with a cagemate in chronic pain condition

**DOI:** 10.1101/2021.12.16.472930

**Authors:** Isabela Miranda Carmona, Paulo Eduardo Carneiro de Oliveira, Daniela Baptista-de-Souza, Azair Canto-de-Souza

## Abstract

The affective component of pain may be shared among conspecifics through emotional contagion, a form of empathic expression. In this sense, reverberation of negative emotions could generate distress behavioral responses, such as pathological anxiety. Evidences reported that amygdala and its benzodiazepine receptors are involved in perception of pain in others. However, relatively little is known about the neural processes underlying emotional contagion induced by pain observation. In the present study, we investigated the effects of midazolam, an allosteric GABAergic receptor agonist, in anxiety-like behaviors induced by cohabitation with cagemate submitted to sciatic nerve constriction. For this purpose, we administrated systemic (0.5, 1.0 and 2.0 mg/kg) and intra-amygdala midazolam injections (3.0 and 30.0 nmol) in observer cagemates before elevated plus-maze (EPM) evaluation. We found that mice subjected to nerve constriction and their observer cagemates increased anxiety-like behavior in the EPM. Further, systemically (1.0 and 2.0 mg/kg) and intra-amygdala administration of midazolam (3.0 and 30 nmol) reverse this anxiogenic effect. Collectively, these results suggest that social interaction with a cagemate under chronic pain produces anxiety-like responses that could be blocked through midazolam application.

## Introduction

Emotional contagion is a rudimentary form of empathy which the observer shares the emotions of another by the perception of emotional or arousal state [1,2]. This affective sharing could be seemed through the pain and its social reverberation. Evidences have shown that pain might be socially perceived between conspecifics and in some cases should provoke dysfunctional behavioral consequences, such as anxiety, depression, cognition impairment and pain-related sensibility disorders [3,4]. From clinical point of view, these disturbances are recognized in family members and/or caregivers of patients in chronic illness conditions. In this context, researches demonstrated elevated psychological distress, anxiety and depression symptoms in spouses of individuals with chronic pain [5,6]. Likewise, in several works, caregivers reported a pronounced unpleasant emotional experience elicited by the burden of providing care and the perception of patients suffering, which contributes to the emergence of anxiety and depressive mood [4,7–9]. The pain-related emotional contagion is possible due its affective component which encodes the disagreeable aspects of pain [10,11].

Given the clinical relevance of these deteriorative mental occurrences, few researches have investigated the neurobiological mechanisms underlying these processes. Therefore, rodent models of emotional contagion employing a conspecific under physical harm might provide efficient evidences about emotional dysfunction observed in caregivers. Among some paradigms to investigate the effects pain-related emotional contagion [12–14], our research group adopted a preclinical approach which cagemates are co-housed with mice submitted to sciatic nerve injury [15–20]. Our previous findings demonstrated that observer/witness cagemates exhibit increased anxiety-like behaviors in the EPM [15,17,18] and hyperalgesia in writhing test [15,16,19,20].

Neuroimaging researches reported that observing someone in pain may trigger an affective/cognitive state which recruits neural structures engaged in experience of empathy [21,22]. Benuzzi and co-workers [23], for example, employing functional magnetic resonance imaging (fMRI), observed higher bilaterally activation of amygdala in volunteers watching videos exhibiting facial expressions of pain compared to non-painful expressions. Interestingly, in studies conducted in youths with psychopathic traits displaying disruptive empathic pain sensitivity, found reduced activation of the amygdala during the presentation of pictures featuring pain situations [24]. Furthermore, pre-clinical approaches have demonstrated the involvement of the amygdala in empathy-related behaviors. For instance, studies showed increased amygdala activation of rats in an emotional contagion model, which cagemates express emotional arousal after receiving footshocks [25,26]. The amygdala is also required during the exhibition of empathic behaviors, such as prosocial activity [27] and observational fear learning [28]. Thus, studies have suggested a fundamental role of the amygdala in attribution of emotions to others [29].

The amygdala is a key structure in the manifestation of the affective component of pain [10,11] and it has been demonstrated that GABA_A_ receptors (γ-aminobutyric acid subtype A receptor) inside the amygdala could be implicated in affective process of pain in rats under neuropathic pain condition [30]. Regarding the high density of GABA_A_ receptors within the amygdala [31], midazolam (an allosteric agonist of GABA_A_ receptor, substance belonging to the group of benzodiazepine drugs) was employed in the assessment of anxiety-like behaviors provoked by chronic pain condition. In this context, some evidence indicated that midazolam reversed anxiogenic-like behavior of rats tested on the EPM [32,33]. In spite of the growing number of studies focusing on neural mechanisms of empathy, no research has examined the effects of benzodiazepines in emotional contagion induced by cohabitation with conspecific submitted to chronic pain. Using an interesting paradigm, Ben-Ami Bartal and colleagues [34] observed that midazolam-treated rats diminished the door-opening of a restrainer to free a trapped cagemate. In this work they argued that midazolam impaired the affective processes necessary for the occurrence of prosocial/helping behavior in rats, suggesting a midazolam effect that goes beyond the anxiolysis.

Considering the social reverberation of pain from one mouse to another and the involvement of amygdala in perception of an affective arousal demonstrated by a conspecific, we proposed to investigate the effect of midazolam over anxiety induced by emotional contagion. For this purpose, we carried out three experiments: (1) evaluation of anxiety-like behavior in mice submitted to chronic pain; (2) effect of systemic and (3) intra-amygdala midazolam administration in the anxiety-like behavior displayed by mice living with a conspecific in chronic pain condition.

## 2. Methods

### 2.1. Subjects and ethics

For this study 382 male Swiss mice (18-20 g) obtained from the animal breeding facility of the Federal University of São Carlos, São Paulo, Brazil, were moved to the animal facility of Psychobiology Group laboratory. After one week for habituation, mice were housed in two per cage [19 cm (width) x 30 cm (length) x 14 cm height)]. Mice were maintained under regular light-dark cycle (12h/12h, lights on at 7:00 a.m.) and controlled temperature (24 ± 1°C) with unrestricted access to food and water except during the brief test periods. The experiments were carried out during the light phase. All procedures were in accordance with the recommended protocol approved by Brazilian Guidelines for Care and Use of Animals for Scientific and Educational Purposes, elaborated by The National Council of Control of Animal Testing (CONCEA). This study was approved by Ethics Committee on Animal Experiments (CEUA 7315090415).

### 2.2. Drugs

Midazolam [allosteric GABA_A_ receptor whose pharmacological activity consists in the potentiation of inhibitory effects of GABA neurotransmitter through the ligation of benzodiazepine site (Roche, Brazil)] was dissolved in saline (0.9% NaCl), injected intraperitoneally at doses of 0.5, 1.0 and 2.0 mg/kg for systemic experiment and microinjected at doses of 3.0 and 30 nmol/0.1μl, for intra-amygdala study. The doses were based on previous works [20,35].

### 2.3. Sciatic nerve constriction

The pain chronic model procedure was based on previous studies [15,18,20,36]. Mice were anesthetized under ketamine and xylazine (100 and 10 mg/kg, i.p., respectively); a linear skin incision was made along the lateral surface of the biceps femoris of right hindpaw and forceps were inserted into the core of the muscle to split the muscle fibers and expose the sciatic nerve. The tissue around the nerve was carefully cut at a distance of approximately 8 mm and then the nerve was compressed with four ligatures using sterile non-inflammatory mononylon 6.0 threads (NC group). For mice in the SHAM group, the sciatic nerve was exposed, according to the procedure described above, however it was not constricted.

### 2.4. Hot plate test

The animals were placed on a heated plate maintained at 52°C and the time to hindpaw shake or hindpaw lick was considered latency of hindpaw withdrawal, measured in seconds, which the constriction injury was performed or not. To avoid injury in the paws, the animals remained on the plate for 30s as a maximum time [15,20].

### 2.5. Elevated plus maze (EPM)

The apparatus was similar to that described in previous studies [37] and comprised two open arms (30 × 5 × 0.25 cm) and two closed arms by transparent glass (clear walls) (30 × 5 × 15 cm) joined to a common central platform (5 × 5 cm). The apparatus was constructed from wood (floor) and was raised to a height of 38.5 cm above floor level. Each mouse was placed on the center of the maze facing an open arm, five minutes after intra-amygdala microinjection. Conventional measures comprised percentage of open arm entries [(open/total entries) x 100], percentage of time spent in open arms [(open/total time) x 100] and number of closed arm entries. Complementary behaviors were percentage of time spent in central platform [(central/total time) x 100], total number and percentage of protected head-dipping [(protected/total) x 100] (HD: exploratory movement of head/shoulders over side of the maze) as well a total number and percentage of stretched attend postures [(protected/total) x 100] (SAP: exploratory posture in which the mouse stretches forward and retracts to original position without any forward locomotion). The closed arms and the central platform were together designated “protected” areas (i.e., offering relative security), while the open arms were designated “unprotected” areas [38]. The duration of test sessions were 5min and, among each test, the apparatus was cleaned with ethanol 20% and dried clothes. All sessions were recorded by a vertically mounted camera linked to a computer for posterior analysis and conducted under moderate illumination (77 lux; measured on the central platform of the EPM) during the light phase of the light-dark cycle. Test videos were scored by a highly trained observer using the free software X-PloRat [39].

### 2.6. Surgery and microinjection

For experiment 3, bilateral stainless-steel guide cannulae (25-gauge x 7 mm long, Insight Instruments, Brazil) were implanted in mice anesthetized by intraperitoneally injection of ketamine hydrochloride (100 mg/kg) plus xylazine (10 mg/kg). Stereotaxic coordinates to target the amygdala were 1.1 mm posterior to bregma, ± 3.3 mm lateral and 3.0 mm ventral from the skull surface [40]. The guide cannulae were fixed to the skull with dental acrylic and jeweler’s screw. Dummy cannulae (33-gauge stainless steel wire; Fishtex®, Brazil) was inserted into the guide cannulae to reduce the incidence of occlusion. At the end of the stereotaxic surgery, the animals received intraperitoneally injections of anti-inflammatory ketoprofen (5 mg/kg) and antibiotic ceftriaxone (4 mg/kg) [41,42]. Before tests mice remained, at least, for 4 to 5 days to recover from the surgery [41]. Midazolam or saline were injected into the amygdala by microinjections units (33-gauge stainless steel cannula; Insight Instruments, Brazil), which extended 1.0 mm beyond the tips of the guide cannula. Each microinjection unit was attached to a 10 μl Hamilton microsyringe via polyethylene tube (PE-10), and administration was programmed by an infusion pump (Insight BI2000, Insight Instruments, Brazil) to deliver a volume at rate of 0.1 μl over a period of 60s. The microinjection procedure consisted of gently restraining the animal, removing the dummy cannula, inserting the injection unit and proceeding with the infusing the solution. The movement of a small air bubble in the polyethylene tube before, during and after the microinjections confirmed the flow of the solution [18,20,35].

### 2.7. Histological procedure

Mice were anesthetized at the end of experiment 3 with ketamine hydrochloride and xylazine solution (100 mg/kg and 10 mg/kg, i.p., respectively) and received a bilateral injection of 0.1 μl solution of 2% of Evans blue, in accordance to the procedure described for intra-amygdala injections. After infusion, animals were euthanized in a CO_2_ chamber; their brains were removed and accommodated in containers containing formalin solution (10%). Subsequently, the brains were sectioned (40 μm) in a cryostat at −20°C (LEICA CM 1850, Leica Biosystems, Germany) for histological analysis of the injection sites. Coronal sections were analyzed by microscope (Olympus BX41, Olympus, Japan) and the visualization of the Evans blue dispersion indicated the sites of the microinjections [40]. Data from mice with injection sites outside the amygdala were excluded from analysis.

### 2.8. General procedure

#### 2.8.1. Experiment 1. Evaluation of anxiety-like behavior in mice submitted to chronic pain

At the weaning [1^st^ experiment day (1^st^ day) or post-natal day 21 (PND21)], mice were housed in pairs and left undisturbed for 14 days until PND35 (14^th^ day), except for cage cleaning [15,18,20,36]. This 14 days period was adopted for development of familiarity between the dyads, based on previous data published by Langford and co-workers [3]. At the 15^th^ day (PND36) one mouse from the pairs was submitted to sciatic nerve constriction (NC) or just surgery procedure without nerve injury (Sham). Mice returned to homecage and remained together for more 14 days (28^th^ day or PND50). At the 28^th^ day, NC and Sham mice were tested in the EPM for assessment of anxiety-like behavior (Figure 1A).

**Figure 1.**
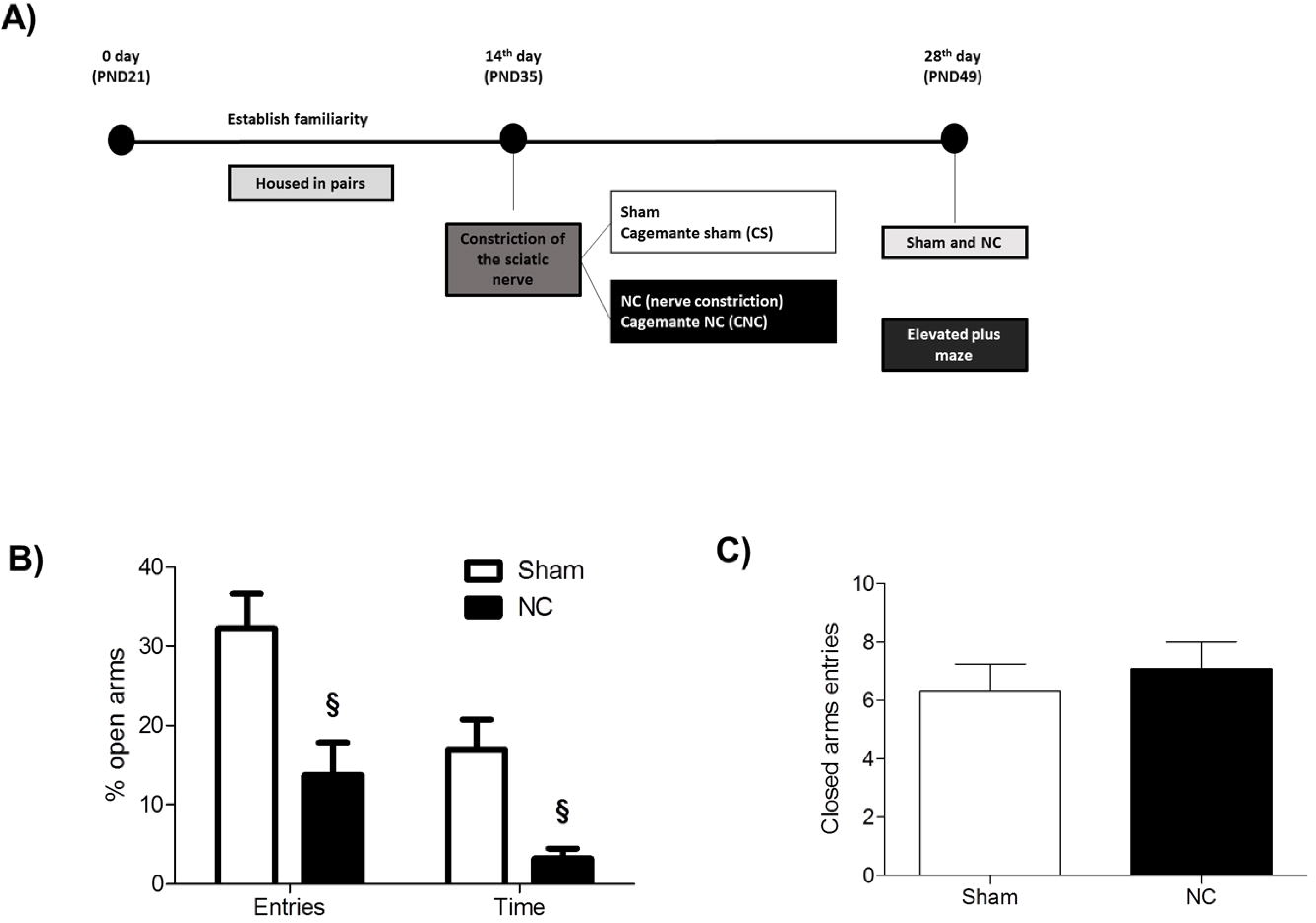
All data are presented as mean ± SEM. (A) Schematic representation of the experimental protocol; (B) effects of chronic pain in percentage of entries and time spent in open arms in mice submitted to nerve constriction and exposed to the EPM test; (C) effects of chronic pain in closed arms entries in mice submitted to nerve constriction and tested in the EPM (n=10 per group). §p < 0.05 vs. sham. Student’s *t* test. NC, nerve constriction.

#### 2.8.2. Experiment 2. Effect of systemic midazolam injection on the anxiety-like behavior in mice living with a conspecific in chronic pain condition

Mice housed with NC or Sham groups were tested in the experiment 2 for evaluation of anxiety-like behavior induced by cohabitation with a cagemate in chronic pain condition. Thus, mice that were not submitted to nerve constriction or sham procedure (described in experiment 1) were named cagemate NC (CNC) or cagemate Sham (CS) starting at the 15^th^ day. At the 28^th^ day, mice from CNC and CS groups received an intraperitoneally injection of saline or midazolam in one of the three doses, 0.5, 1.0 or 2.0 mg/kg. Thirty minutes after drug administration, mice were tested in the EPM for assessment of anxiety-like behavior. The NC and Sham animals used in this experiment were placed in the hot plate for evaluation of surgery effectiveness (Figure 2A).

**Figure 2.**
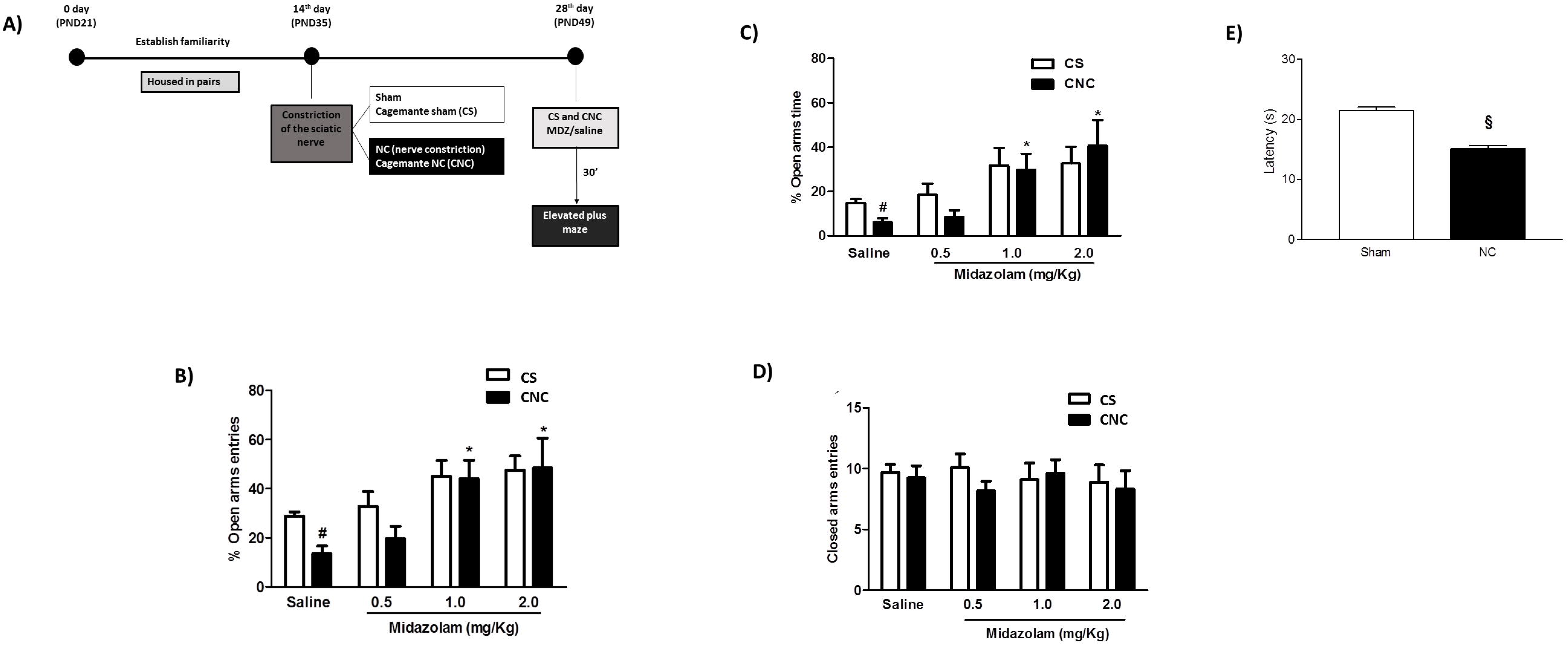
All data are presented as mean ± SEM. (A) Schematic representation of the experimental protocol; (B) anxiolytic effect of systemic midazolam (0.5, 1.0 e 2.0 mg/kg, i.p) in percentage of entries and (C) time spent in open arms of the EPM in mice that co-housed with conspecific under chronic pain; (D) effects of midazolam (0.5, 1.0 e 2.0 mg/kg, i.p) injection in closed arms entries of EPM in mice submitted to cohabitation with conspecific in chronic pain condition (n=9-12 per group). #p<0.05 vs. respective CS group. *p<0.05 vs. respective saline group. Two-way ANOVA followed by Duncan’s post-hoc test. CS, cagemate Sham; CNC, cagemate nerve constriction. (E) Chronic pain induces thermal hyperalgesic behaviors in mice submitted to sciatic nerve constriction (n=51 per group). §p < 0.05 vs. sham. Student’s *t* test. NC, nerve constriction.

#### 2.8.3. Experiment 3. Effect of midazolam microinjected intra-amygdala on the anxiety-like behavior in mice living with a conspecific in chronic pain condition

Other subset of mice were submitted to the same protocol described in Experiment 2 (CNC and CS groups) except that in the 23^th^ day bilateral cannulae were surgically implanted in the amygdala of the cagemate. At the 28^th^ day, subjects from CNC and CS groups received intra-amygdala saline or midazolam (3.0 or 30.0 nmol/0.1μl) microinjections 5 minutes before the EPM test. The NC and Sham animals used in this experiment were placed in the hot plate for evaluation of surgery effectiveness (Figure 3A).

**Figure 3.**
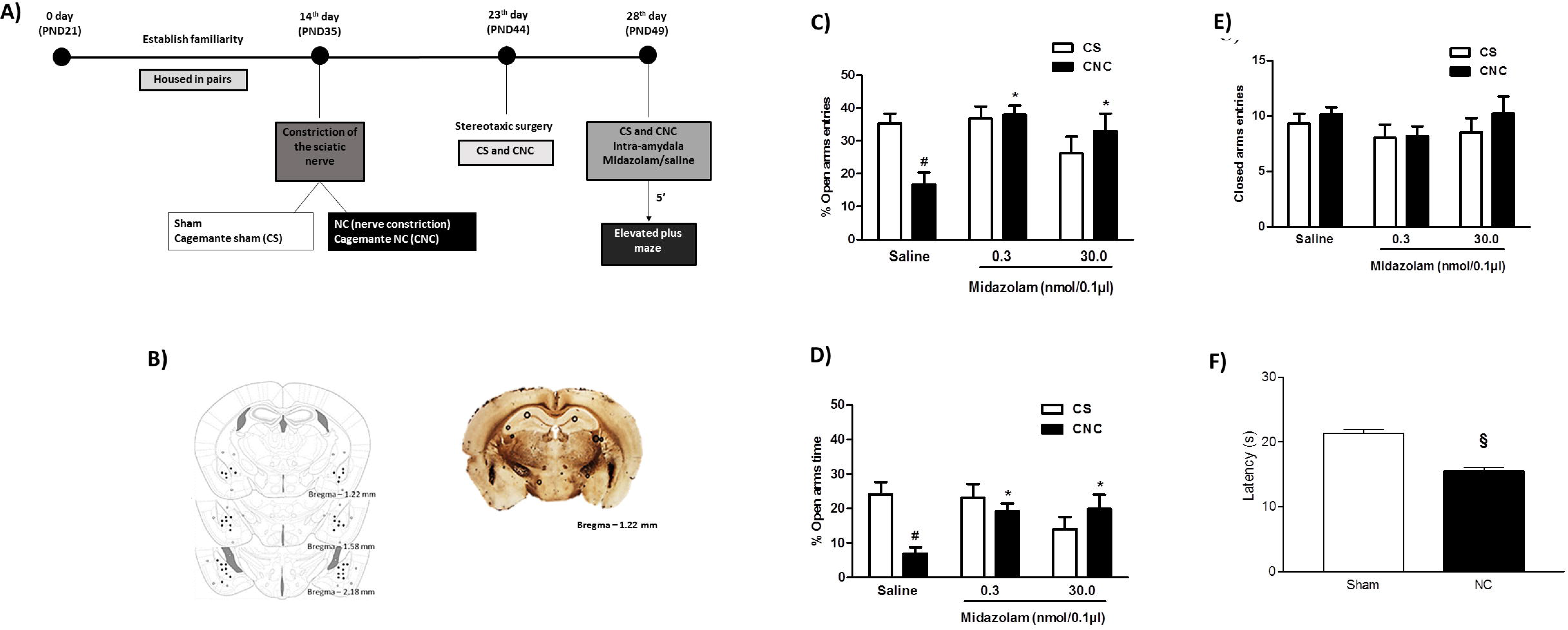
All data are presented as mean ± SEM. (A) Schematic representation of the experimental protocol; (B) schematic representation of microinfusion sites within (black circles) or outside (gray circles) the amygdala. Sections are between −1.22 mm and −2.18mm from the bregma. The points in the illustration represent some of the sites reached. Photomicrography of a coronal section (Bregma - 1.22 mm) of a representative subject showing an injection site within the mouse amygdala; (C) effects of midazolam (3.0 and 30.0 nmol/0.1μl) intra-amygdala injection in percentage of entries and (D) time spent in open arms in mice submitted to cohabitation with conspecific in chronic pain and exposed to elevated plus-maze; (E) effects of midazolam (3.0 and 30.0 nmol/0.1μl) intra-amygdala injection in closed arms entries in mice submitted to cohabitation with conspecific under chronic pain and submitted to elevated plus-maze (11-12 per group). #p<0.05 vs. respective CS group. *p <0.05 vs. respective saline group. Two-way ANOVA followed by Duncan’s post-hoc test. CS, cagemate Sham; CNC, cagemate nerve constriction. (F) Chronic pain induces thermal hyperalgesic behaviors in mice submitted to sciatic nerve constriction (n=42-43 per group). §p < 0.05 vs. sham. Student’s *t* test. NC, nerve constriction.

### 2.9. Statistical analysis

All data were expressed as means ± SEM. Data from experiment 1 were analyzed by Student’s *t* test. For experiment 2 and 3 we applied two-way analysis of variance (ANOVA) (cohabitation type x treatment). When ANOVA indicated interaction effects we applied Duncan’s post-hoc test. Statistical differences were considered significant when p value was less than 0.05 (p<0.05).

## 3. Results

### 3.1. Experiment 1. Chronic pain induces increased anxiety behaviors

Twenty were tested in this experiment [Sham (n=10 and NC (n=10)]. Student’s *t* test revealed decrease in percentage of open arms entries [t_(18)_= 3.09; p<0.05] and time spent in open arms [t_(18)_= 3.45; p<0.05] (Figure 1B) demonstrating an anxiogenic-like effect induced by chronic pain condition in mice tested in EPM. The chronic pain did not modify the number of closed arm entries [t_(18)_= −0.61; p=0.55] (Figure 1C) suggesting no locomotor activity interference in animals submitted to nerve constriction.

Student’s *t* test analysis showed significant effect in percentage of protected head-dipping and percentage of protected SAP. No significant effect was observed in percentage of time spent in the center platform, total head-dipping and total SAP (Table 1).

**Table 1.** Complementary behaviors evaluated in the EPM test (experiment 1).

| Behavior | Pain condition |  | Student's *t* test |
| --- | --- | --- | --- |
|  | Sham | NC |  |
| Center time (%) | 41.8 ± 6.6 | 43.4 ± 7.0 | $t_{(18)} = -0.16$; $p=0.87$ |
| Total head-dipping | 28.8 ± 2.3 | 24.8 ± 2.8 | $t_{(18)} = 0.92$; $p=0.37$ |
| Protected head-dipping (%) | 75.5 ± 5.9 | 97.5 ± 1.1§ | $t_{(18)} = -3.70$; $p<0.05$ |
| Total SAP | 40.9 ± 2.7 | 41.4 ± 2.8 | $t_{(18)} = -0.13$; $p=0.90$ |
| Protected SAP (%) | 87.5 ± 2.0 | 97.3 ± 0.9§ | $t_{(18)} = -4.36$; $p<0.05$ |

Data are presented as mean + SEM. § $p<0.05$ vs. Sham group. Student's *t* test. CS: cagemate Sham; CNC: cagemate nerve constriction. SAP: stretched attend posture. a: cohabitation type factor; b: treatment factor; c: interaction by cohabitation type and treatment factors.

### 3.2. Experiment 2. Anxiolytic effect of systemic midazolam in the EPM responses exhibited by mice living with a pair in chronic pain condition

Eighty five cagemates were tested in the EPM [CS/Sal (n=12), CNC/Sal (n=11), CS/Mdz 0.5 (n=11), CNC/Mdz 0.5 (n=12), CS/Mdz 1.0 (n=10), CNC/Mdz 1.0 (n=11), CS/Mdz 2.0 (n=9) and CNC/Mdz 2.0 (n=9)]. For percentage of open arms entries (Figure 2B), two-way ANOVA revealed effects of cohabitation type [F_(1,77)_=7.94; p<0.05] and treatment [F_(3,77)_=5.40; p<0.05] factors, but no interaction between them [F_(3,77)_=1.33; p=0.27]. Two-way ANOVA also demonstrated effects of cohabitation type [F_(1,77)_=7.26; p<0.05] and treatment [F_(3,77)_=6.48; p<0.05), but not significant effect of interaction between the factors [F_(3,77)_=1.24; p=0.30] in time spent in open arms (Figure 2C). Duncan’s test showed that the mice housed with animals submitted to chronic pain injury (CNC) decreased the open arms exploration compared to cagemate Sham (CS), indicating an anxiogenic-like effect induced by cohabitation with a pair in pain condition (Figures 2B and 2C). ANOVA followed by Duncan’s test indicated that the midazolam treatment (1.0 and 2.0 mg/kg) increased the percentage of open arm entries and percentage of time spent in open arms in CNC animals, compared to respective saline group (Figures 2B and 2C). There are no effects of cohabitation type [F_(1,77)_=0.56; p=0.46], treatment [F_(3,77)_=0.22; p=0.88] and interaction [F_(3,77)_=0.44; p=0.72] in closed arms entries, indicating no locomotor impairment in all tested groups (Figure 2D). One hundred and two mice were tested in the hot plate [Sham (n=51) and NC (n=51)]. Results revealed that the latency of response to the thermal stimulus was significantly reduced in animals submitted to sciatic nerve constriction compared to Sham group [t_(100)_= 9.04, *p*<0.05] (Figure 2E).

Statistical analysis showed that midazolam treatment (1.0 and 2.0 mg/kg) increased percentage of protected head-dipping in both CS and CNC animals compared to respective saline group. The higher dose (2.0 mg/kg) increased total head-dipping and percentage of protected SAP in CNC animals compared to respective saline group (Table 2).

**Table 2.** Complementary behaviors evaluated in the EPM test (experiment 2).

| Behavior | Midazolam (mg/Kg) |  |  |  |  |  |  |  | Two-way ANOVA |
| --- | --- | --- | --- | --- | --- | --- | --- | --- | --- |
|  | Saline |  | 0.5 |  | 1.0 |  | 2.0 |  |  |
|  | CS | CNC | CS | CNC | CS | CNC | CS | CNC |  |
| Center time (%) | 24.5 ± 3.0 | 25.4 ± 3.7 | 24.2 ± 2.0 | 21.4 ± 2.8 | 22.2 ± 2.4 | 24.2 ± 3.0 | 28.2 ± 5.5 | 16.5 ± 1.1 | a $F_{(1,77)}=1.77$; $p=0.19$<br>b $F_{(3,77)}=0.28$; $p=0.84$<br>c $F_{(3,77)}=1.73$; $p=0.17$ |
| Total head-dipping | 31.1 ± 3.5 | 29.4 ± 3.3 | 29.0 ± 1.6 | 25.0 ± 2.7 | 36.2 ± 3.4 | 43.7 ± 6.4 | 37.6 ± 4.0 | 42.4 ± 6.7* | a $F_{(1,77)}=0.29$; $p=0.59$<br>b $F_{(3,77)}=4.81$; $p<0.05$<br>c $F_{(3,77)}=0.80$; $p=0.50$ |
| Protected head-dipping (%) | 76.3 ± 3.9 | 87.2 ± 4.2 | 61.0 ± 7.5 | 79.1 ± 6.5 | 47.0 ± 7.8* | 52.7 ± 8.3* | 47.1 ± 9.6* | 42.5 ± 13.1* | a $F_{(1,77)}=1.83$; $p=0.18$<br>b $F_{(3,77)}=9.56$; $p<0.05$<br>c $F_{(3,77)}=0.70$; $p=0.55$ |
| Total SAP | 40.6 ± 3.8 | 45.1 ± 5.1 | 48.4 ± 4.7 | 46.0 ± 5.1 | 41.7 ± 4.7 | 44.8 ± 6.8 | 42.2 ± 6.6 | 46.6 ± 7.2 | a $F_{(1,77)}=0.37$; $p=0.54$<br>b $F_{(3,77)}=0.27$; $p=0.85$<br>c $F_{(3,77)}=0.18$; $p=0.91$ |
| Protected SAP (%) | 74.8 ± 4.8 | 89.3 ± 3.5 | 79.2 ± 6.4 | 88.1 ± 3.9 | 70.6 ± 6.3 | 73.0 ± 7.3 | 69.5 ± 6.2 | 60.4 ± 10.4* | a $F_{(1,77)}=0.88$; $p=0.35$<br>b $F_{(3,77)}=3.74$; $p<0.05$<br>c $F_{(3,77)}=1.21$; $p=0.31$ |

Data are presented as mean + SEM. \* $p<0.05$ vs. respective saline group. Two-way ANOVA followed by Duncan's post-hoc test. CS: cagemate Sham; CNC: cagemate nerve constriction. SAP: stretched attend posture. a: cohobitation type factor; b: treatment factor; c: interaction by cohobitation type and treatment factors.

### 3.3. Experiment 3. Anxiolytic effect of intra-amygdala injection of midazolam in the EPM responses exhibited by mice living with a pair in chronic pain condition

The histological analysis confirmed that 68 mice received cannula implantation in the amygdala complex [CS/Sal (n=11), CNC/Saline (n=11), CS/Mdz 3.0 (n=11), CNC/Mdz 3.0 (n=11), CS/Mdz 30.0 (n=12) and CNC/Mdz 30.0 (n=12)] (Figure 3B). Two-way ANOVA revealed significant effects in percentage of open arm entries for interaction between cohabitation type and treatment [F_(2,62)_=5.41; p<0.05], treatment [F_(2,62)_=3.92; p<0.05), however no significant effect for cohabitation type [F_(1,62)_=1.21; p=0.28] (Figure 3C). This analyses also demonstrated significant effect in cohabitation type x treatment for percentage of time spent in open arms [F_(2,62)_=5.49; p<0.05], without significant effects on cohabitation type [F_(1,62)_=3.19; p=0.08] and treatment [F_(2,62)_=1.50; p=0.23] (Figure 3D). Post hoc analyses revealed that the group CNC decreases the open arms exploration, an anxiogenic-like effect, compared to CS. These effects were attenuated by midazolam intra-amygdala injections (3.0 and 30.0 nmol/0.1μl) represented by increase in percentage of open arms entries and percentage of time spent in open arms (Figures 3C and 3D). Living with a partner in chronic pain condition [F_(1,62)_=1.04; p=0.31] and midazolam administration [F_(2,62)_=1.08; p=0.34] did not modify the number of closed arm entries. Also, there is no interaction between cohabitation type and treatment factors [F_(2,62)_=0.23; p=0.79], suggesting no locomotor activity impairment in all groups (Figure 3E). Eighty nine mice were tested in the hot plate [Sham (n=44) and NC (n=45)]. Results indicated a decrease in the latency of hindpaw withdrawal after thermal stimulus in NC group compared to Sham animals [t_(87)_ = 7.02, p<0.05] (Figure 3F).

Two-way ANOVA followed by Duncan’s test revealed statistically significant effects in CNC animals in percentage of protected head-dipping, protected SAP and percentage of time spent in the center platform compared to CS. Midazolam treatment (3.0 and 30.0 nmol/0.1μl) reversed the influence of cohabitation type in percentage of protected head-dipping, protected SAP, percentage of time spent in the center platform and the higher dose (30.0 nmol/0.1μl) also increased total SAP in CNC animals compared to respective saline group (Table 3).

**Table 3.**
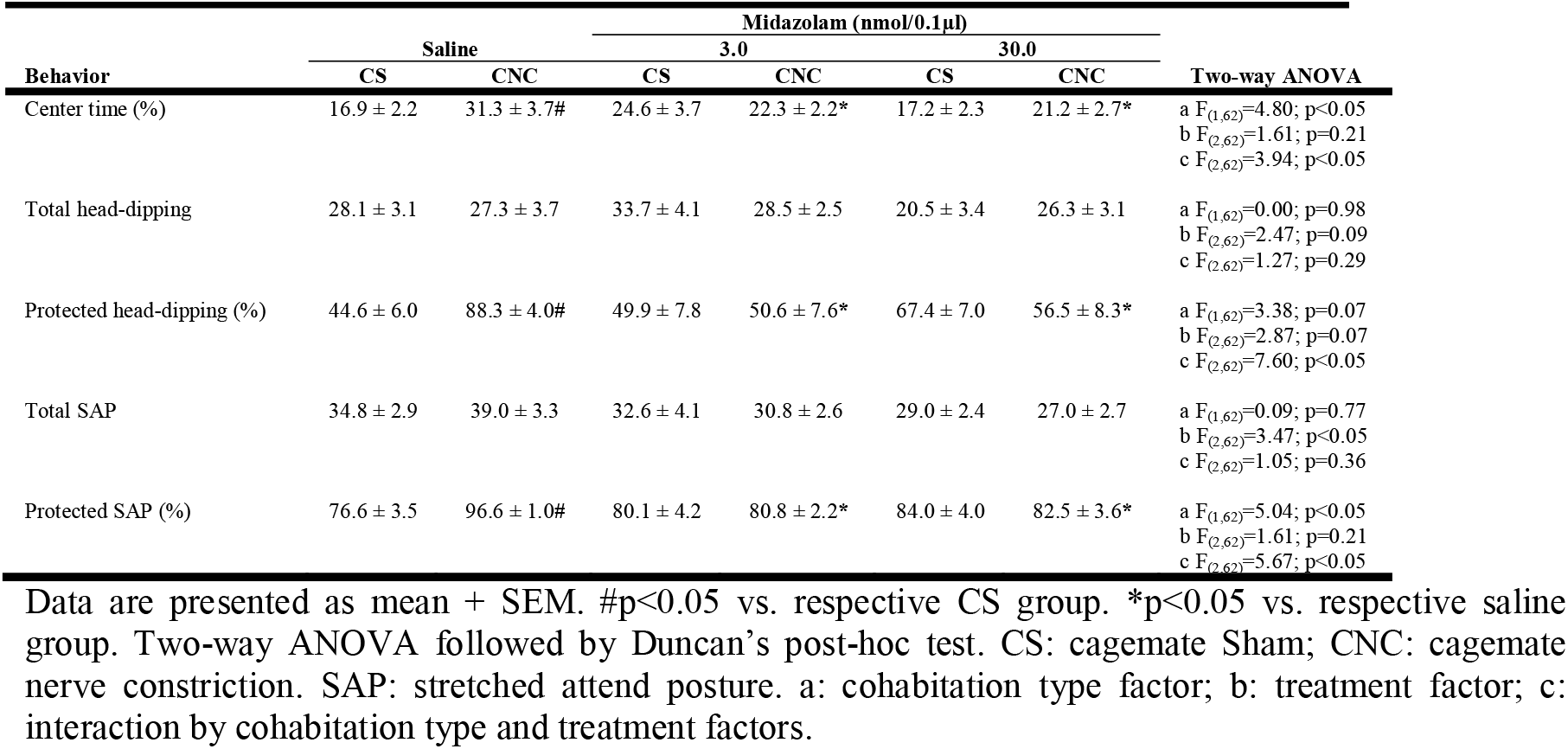
Complementary behaviors evaluated in the EPM test (experiment 3).

| Behavior | Midazolam (nmol/0.1μl) |  |  |  |  |  | Two-way ANOVA |
| --- | --- | --- | --- | --- | --- | --- | --- |
|  | Saline |  | 3.0 |  | 30.0 |  |  |
|  | CS | CNC | CS | CNC | CS | CNC |  |
| Center time (%) | 16.9 ± 2.2 | 31.3 ± 3.7# | 24.6 ± 3.7 | 22.3 ± 2.2* | 17.2 ± 2.3 | 21.2 ± 2.7* | a F<sub>(1,62)</sub>=4.80; p<0.05<br>b F<sub>(2,62)</sub>=1.61; p=0.21<br>c F<sub>(2,62)</sub>=3.94; p<0.05 |
| Total head-dipping | 28.1 ± 3.1 | 27.3 ± 3.7 | 33.7 ± 4.1 | 28.5 ± 2.5 | 20.5 ± 3.4 | 26.3 ± 3.1 | a F<sub>(1,62)</sub>=0.00; p=0.98<br>b F<sub>(2,62)</sub>=2.47; p=0.09<br>c F<sub>(2,62)</sub>=1.27; p=0.29 |
| Protected head-dipping (%) | 44.6 ± 6.0 | 88.3 ± 4.0# | 49.9 ± 7.8 | 50.6 ± 7.6* | 67.4 ± 7.0 | 56.5 ± 8.3* | a F<sub>(1,62)</sub>=3.38; p=0.07<br>b F<sub>(2,62)</sub>=2.87; p=0.07<br>c F<sub>(2,62)</sub>=7.60; p<0.05 |
| Total SAP | 34.8 ± 2.9 | 39.0 ± 3.3 | 32.6 ± 4.1 | 30.8 ± 2.6 | 29.0 ± 2.4 | 27.0 ± 2.7 | a F<sub>(1,62)</sub>=0.09; p=0.77<br>b F<sub>(2,62)</sub>=3.47; p<0.05<br>c F<sub>(2,62)</sub>=1.05; p=0.36 |
| Protected SAP (%) | 76.6 ± 3.5 | 96.6 ± 1.0# | 80.1 ± 4.2 | 80.8 ± 2.2* | 84.0 ± 4.0 | 82.5 ± 3.6* | a F<sub>(1,62)</sub>=5.04; p<0.05<br>b F<sub>(2,62)</sub>=1.61; p=0.21<br>c F<sub>(2,62)</sub>=5.67; p<0.05 |

Data are presented as mean + SEM. # $p<0.05$ vs. respective CS group. \* $p<0.05$ vs. respective saline group. Two-way ANOVA followed by Duncan's post-hoc test. CS: cagemate Sham; CNC: cagemate nerve constriction. SAP: stretched attend posture. a: cohobitation type factor; b: treatment factor; c: interaction by cohobitation type and treatment factors.

## 4. Discussion

The present study investigated whether cohabitation with a conspecific in chronic pain condition would provoke changes in anxiety-like behavior in cagemates and whether systemic or intra-amygdala administration of midazolam would be able to reverse this effect of pain-induced emotional contagion. Confirming our previous results, we found that living, for 14 days, with mice submitted to sciatic nerve constriction induces anxiogenic-like behavior in cagemates tested in the EPM [15,17,18] and midazolam reversed this anxiogenic effect by both, systemic and intra-amygdala administration. In addition to data showing midazolam effects on the EPM test, we complimented these previous findings showing increase in anxiety-like behaviors in mice subjected to neuropathic pain condition when compared to Sham group.

Firstly, we have to highlight the importance of results found in experiment 1. Considering we are studying the neural basis of emotional contagion, the assessment of anxiety-like behavior in NC group is essential to show that mice submitted to nerve constriction are sharing not only the discomfort of chronic pain, but also a negative emotional state. Our group had already observed stress-induced emotional contagion through the increase in anxiety-like behavior in both, chronic restrained mice and their cagemates [36].

The anxiogenic effect of sciatic nerve constriction in NC group suggests that our 14 day in pain condition protocol is able to induce anxiety in this group. Furthermore, NC group exhibited ipsilateral thermal hyperalgesia, compared to Sham mice, observed by decrease in latency of hindpaw withdrawal in the hot plate paradigm. It confirms that NC mice were still in pain condition until the day of the EPM test. Thus, these findings corroborate some researches which investigated the emotional consequences of chronic neuropathic pain. Several studies demonstrated that mice submitted to neuropathic pain condition showed heat thermal hyperalgesia, increase in mechanical allodynia and anxiogenic-like behaviors at least for 4 weeks after the surgery [43–49].

The emotional contagion induced by distress situation in a conspecific has been explored by our group using chronic sciatic nerve constriction [15,17,18] or chronic restraint stress [34]. These studies observed that cohabitation with a mouse submitted to chronic pain or chronic stress promotes decrease in exploration of open arms of the EPM. Our findings are supported by works verifying the emotional contagion through protocols which the cagemate witness or coexists with aversive stimuli. For instance, Tomiyoshi and collaborators [14] observed that cagemates co-housed, for 20 days, with mice melanoma-bearer reduced number of entries and time spent in open arms, compared to health partner. In addition, still considering the pain as an aversive stimulus, studies showed anxiogenic-like behaviors in cagemates that cohabitated with rats submitted to chronic inflammatory pain [12,13]. In this context, several researches reported increase in anxiety-like behavior exhibited by cagemates after witnessing their partners suffering repeated defeat stress [50–53]. Else, Yang and colleagues [54] demonstrated enhanced avoidance of open arms by cagemates living with mice models of brain disorders (epilepsy or depression) in EPM test. Importantly, some of these works provided evidence emphasizing that unstable psychosocial environments elicit paralleled behaviors in physically harm and vicariously distressed mice [12,36,51,53,54], supporting our data reporting diminished open arms exploration in both, NC and CNC groups. Taken together, these results suggest that emotional states mighty affect the subject and object bidirectionally [1,3,22].

In this context, although benzodiazepines are frequently indicated for clinical treatment of pain-induced anxiety, their clinical use with analgesic intent is still controversial [55,56]. Here, we found that systemic injection of midazolam brought back the anxiety behavior of CNC group to CS levels. Interestingly, this is the first evidence of midazolam effects in vicarious pain-induced anxiety behavior. However, our results are in agreement previous studies showing that systemic midazolam or diazepam reversed anxiogenic-like behavior, induced by different models of chronic pain, such as orofacial cancer, constrict nerve injury, persistent inflammation and neuropathy [32,33,57,58]. Moreover, in other approaches, intraperitoneally midazolam also reversed distress-induced anxiety and fear. For instance, midazolam restored the anxiety- and fear-responses to control group levels in rats submitted to stressful, pharmacological- or contextual-aversive stimuli [59–62].

Due to high density of GABA_A_ receptors in the amygdala [31] and its role in emotional recognition [29], we investigated the effects of bilateral microinjection of midazolam in pain-induced emotional contagion. Thus, we found that intra-amygdala administration of midazolam also provoked anxiolytic-like behaviors in cagemates co-housed with a partner submitted to chronic pain. An extensive body of evidence has shown anxiolytic-like behaviors in rodents through microinjection of benzodiazepines within the amygdala. In different studies, intra-amygdala administration of midazolam or diazepam reduced anxiety- and fear-related behaviors, such as diminished time of defensive freezing behavior preceded, or not, by restraint stress [60,63,64], reduced time spent exhibiting electrified probe burying behavior [65], enhanced time spent in central zone of the open field test [66] and augmented open arms exploration of mice, or rats, tested in the EPM on the first or second trials [35,67,68].

Strikingly, systemic and intra-amygdala administration of midazolam promotes its effects only in mice housed with a partner submitted to chronic pain (CNC), but not in cagemates that lived with sham group (CS). Although unexpected, some works had already observed this selective effect of midazolam and despite of these studies did not highlight this effect, administration of midazolam affected only distressed groups. For instance, Bignante and colleagues [59] showed that midazolam reversed the acute stress-induced anxiogenic-like behavior in the EPM test compared to vehicle, whilst midazolam had no effect in non-stressed group compared to saline. In addition, midazolam reduced the freezing duration in low-anxiety rats re-exposured to the context associated to fear conditioning compared to the high-anxiety group [62]. The same research group also observed that midazolam reversed the diminished frequency of entries in the central area of the open field arena and contextual-induced freezing time only in high-anxiety rats chronically treated with corticosterone when compared to vehicle group [61]. In the same way, systemic and local injection of midazolam prevented augmented freezing behavior in fear conditioning test only in rats exposed to restraint stress, showing no effects in non-stressed group [60,63]. Evidences have pointed that benzodiazepines induce their anxiolytic action mainly through GABA_A_ receptor α_2_ subunit, since a mutation on the α_2_ subunit gene blocks the attenuation of anxiety by administration of diazepam [69,70]. Hence, despite being purely speculative, the effects of benzodiazepines in these specifics groups may be consequence of enhanced density of α_2_ subunit. In agreement with this proposition studies have shown that α_2_ protein levels in the amygdala are augmented in subjects underwent to physical, psychological or pharmacological stress [71–74]. Interestingly, our suggestion is reinforced by Jacobson-Pick and co-workers [75] which observed that the increase in α_2_ subunit protein in the amygdala of young stressed rats drives to exclusively anxiolytic effects of diazepam in this group when compared to vehicle. In this context, the initial anxiolytic effects of benzodiazepines would depend on a minimal level of anxiety [75]. Thus, due to the relevance of GABA_A_ receptor α_2_ subunit in promoting the anxiolytic effects of benzodiazepines, future works from our laboratory will objectify this understanding.

In conclusion, the results obtained in this study demonstrated that mice co-housed with cagemates in chronic pain condition exhibited behavioral alterations, likewise the anxiogenic-like responses. Furthermore, systemic and intra-amygdala administration of midazolam was able to reverse this enhanced anxiety behaviors induced by distressing cohabitation. The emotions shared among conspecifics would reflect the concerning and awareness of patient suffering by caregivers and family members which lead to impaired mental health. Thus, our model may be useful to study neural mechanisms emotional contagion and improve the treatment of mental disorders.

## Acknowledgments

The experiments described in this manuscript were funded by CAPES (Coordination for the Improvement of Higher Education Personnel), CNPq (National Council for Scientific and Technical Development) and FAPESP (17/25409–0), Brazil. I.M. Carmona was recipient of CAPES research scholarship; P.E. Carneiro de Oliveira received PNPD/CAPES scholarship; D. Baptista-de-Souza received a FAPESP research scholarship (2016/0004-6); A. Canto-de-Souza was recipient of CNPq research fellowships (309201/2015–2). The authors would like to thank Lara Maria Silveira for her technical assistance.

## Author Contributions

**Isabela Carmona**: Data curation, Formal analysis, Investigation, Project administration, Writing – original draft preparation, Writing – review & editing. **Paulo Carneiro de Oliveira**: Data curation, Formal analysis, Investigation, Writing – original draft preparation, Writing – review & editing. **Daniela Baptista-de-Souza**: Conceptualization, Methodology. **Azair Canto-de-Souza**: Conceptualization, Funding acquisition, Project administration, Writing – review & editing, Supervision. All authors approved the final version to be published.

